# Cryo-EM Structure of the Cyclase Domain and Evaluation of Substrate Channeling in a Bifunctional Class II Terpene Synthase

**DOI:** 10.1101/2025.08.20.671325

**Authors:** Matthew N. Gaynes, Kollin Schultz, Eliott S. Wenger, Trey A. Ronnebaum, Ronen Marmorstein, David W. Christianson

**Affiliations:** Roy and Diana Vagelos Laboratories, Department of Chemistry, University of Pennsylvania, Philadelphia, Pennsylvania, 19104-6323, USA; Abramson Family Cancer Research Institute, Perelman School of Medicine, University of Pennsylvania, Philadelphia, PA 19104, USA; Graduate Group in Biochemistry, Biophysics, and Chemical Biology, Perelman School of Medicine, University of Pennsylvania, Philadelphia, PA 19104, USA; Department of Biochemistry and Biophysics, Perelman School of Medicine, University of Pennsylvania, Philadelphia, PA 19104, USA

## Abstract

Copalyl diphosphate synthase from *Penicillium verruculosum* (PvCPS) is a bifunctional class II terpene synthase containing a prenyltransferase that produces geranylgeranyl diphosphate (GGPP) and a class II cyclase that utilizes GGPP as a substrate to generate the bicyclic diterpene copalyl diphosphate. The various stereoisomers of copalyl diphosphate establish the greater family of labdane natural products, many of which have environmental and medicinal impact. Understanding structure-function relationships in class II diterpene synthases is crucial for guiding protein engineering campaigns aimed at the generation of diverse bicyclic diterpene scaffolds. However, only a limited number of structures are available for class II cyclases from bacteria, plants, and humans, and no structures are available for a class II cyclase from a fungus. Further, bifunctional class II terpene synthases have not been investigated with regard to substrate channeling between the prenyltransferase and the cyclase. Here, we report the 2.9 Å-resolution cryo-EM structure of the 63-kD class II cyclase domain from PvCPS. Comparisons with bacterial and plant copalyl diphosphate synthases reveal conserved residues that likely guide the formation of the bicyclic labdane core, but divergent catalytic dyads that mediate the final deprotonation step of catalysis. Substrate competition experiments reveal preferential GGPP transit from the PvCPS prenyltransferase to the cyclase, even when prepared as separate constructs. These results are consistent with a model in which transient prenyltransferase-cyclase association facilitates substrate channeling due to active site proximity.

## Introduction

Terpenes comprise a ubiquitous class of natural products with an extensive range of applications as flavors, fragrances, pharmaceuticals, insecticides, and next-generation biofuels.^1–6^ Terpene biosynthesis is rooted in primary metabolism: a prenyltransferase catalyzes the condensation of two C_5_ substrates, dimethylallyl diphosphate (DMAPP) and one or more equivalents of isopentenyl diphosphate (IPP), to produce C_10_ geranyl diphosphate (GPP), C_15_ farnesyl diphosphate (FPP), C_20_ geranylgeranyl diphosphate (GGPP), and even longer isoprenoid diphosphates.^7,8^ These linear isoprenoids have diverse biological functions, e.g., in protein prenylation to direct cellular localization, or small-molecule prenylation to generate intermediates in alkaloid or polyketide biosynthesis.^9–12^ However, the most intricate reactions of terpene biosynthesis are found in secondary metabolism, where isoprenoid cyclization reactions yield products containing multiple rings and stereocenters. These reactions are catalyzed by terpene cyclases, enzymes that direct astoundingly complex carbocation-mediated cyclization cascades.^13–15^

A terpene cyclase catalyzes the first committed step in the biosynthesis of a terpenoid natural product and usually belongs to one of two principal enzyme classes based on domain architecture and chemical strategy for initial carbocation formation. A class I cyclase contains an active site in the middle of an α helical bundle designated as the α domain, with two metal binding motifs at the mouth of the active site. The α domain fold of class I terpene cyclases is shared with prenyltransferases, and both utilize a trinuclear metal cluster^16^ to initiate ionization of the substrate diphosphate group to generate an allylic carbocation. A class II cyclase adopts a different α helical fold, denoted as a β domain that is most often associated with a structurally homologous γ domain.^14,17^ The γ domain is inserted between the first and second helices of the β domain and caps the active site in the β domain. The β domain contains an aspartic acid that protonates the terminal C=C bond (or epoxide) in the isoprenoid substrate to generate a tertiary carbocation.^18^ Of note, class II cyclization products retain the substrate diphosphate group, e.g., as observed for *ent*-copalyl diphosphate synthase.^19,20^

Prenyltransferases and terpene cyclases are found individually across all domains of life, but bifunctional terpene synthases containing both enzymes in a single polypeptide chain have been identified mainly in fungal species.^21^ The first such reported bifunctional terpene synthase, fusicoccadiene synthase from *Phomopsis amygdali* (PaFS), contains a prenyltransferase (GGPP synthase) and a class I diterpene cyclase.^22^ The first bifunctional terpene synthases identified containing a prenyltransferase (GGPP synthase) and a class II cyclase were copalyl diphosphate synthases from *Penicillium verruculosum* (Figure 1) and *Penicillium fellutanum* (PvCPS and PfCPS, respectively).^23^ In both class I and class II bifunctional terpene synthases, the prenyltransferase is connected to the cyclase through a flexible linker containing 40–125 residues, with domain architecture represented as α∼α and α∼βγ, respectively.

**Figure 1.**
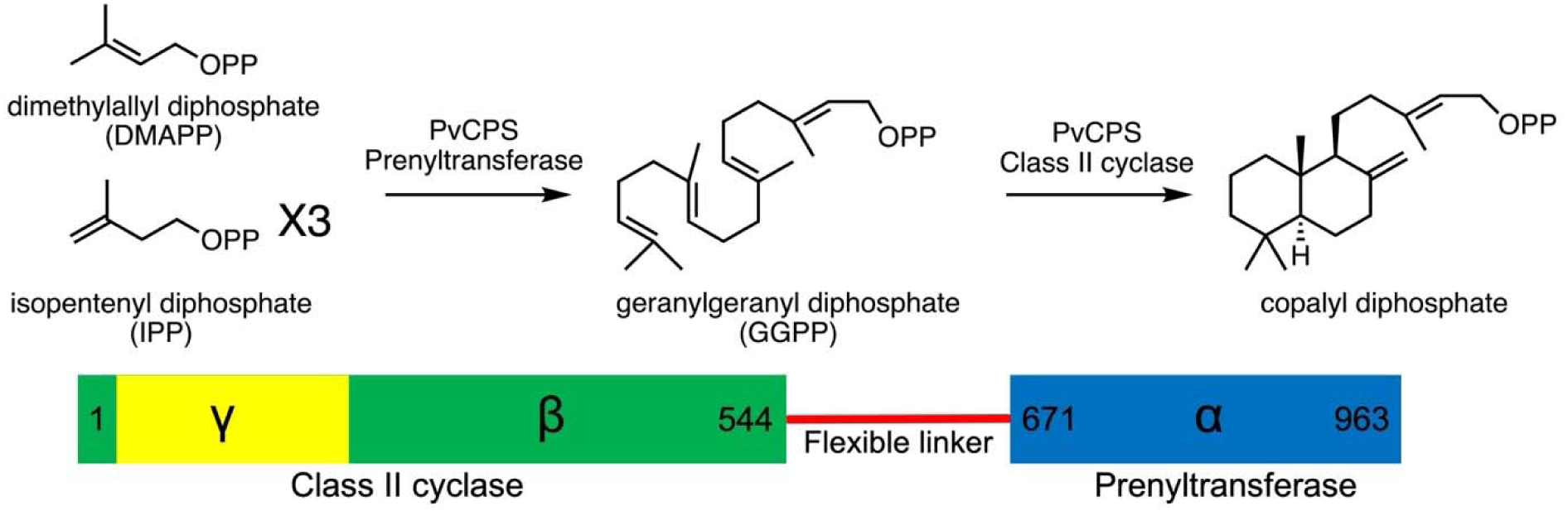
Reaction sequence catalyzed by PvCPS. Prenyltransferase substrates DMAPP and IPP are utilized by the prenyltransferase α domain to produce GGPP. The class II cyclase active site at the βγ domain interface then cyclizes GGPP to generate copalyl diphosphate. The primary structure of PvCPS illustrated at bottom shows that the γ domain is inserted between the first and second helices of the β domain. The γ domain is structurally homologous to the β domain and is thought to be the result of primordial gene duplication and fusion. A 125-residue flexible linker connects the prenyltransferase to the cyclase.

Bifunctional terpene synthases have been referred to as “assembly-line” synthases^24^ because some (but not all) class I bifunctional synthases exhibit substrate channeling in the first two reactions of terpene biosynthesis.^25–27^ Cryo-EM studies of PaFS^28,29^ and the related bifunctional synthase variediene synthase from *Emericella variecolor* (EvVS)^27^ reveal a central prenyltransferase octamer or hexamer, respectively, surrounded by class I cyclase domains with overall (α∼α)_8_ or (α∼α)_6_ architecture. In PaFS, cyclase domains are randomly splayed out around the central prenyltransferase octamer but are capable of transient binding to the side of the prenyltransferase oligomer, which is believed to facilitate intramolecular substrate channeling.^26,28,29^ Intramolecular substrate channeling is not observed in EvVS, and cyclase domains are fixed in apical positions above and below the prenyltransferase hexamer.^27^

To date, substrate channeling has not been explored in any class II bifunctional terpene synthase. Moreover, no high-resolution structure of a bifunctional prenyltransferase-class II cyclase has been reported to date. Even so, the X-ray crystal structure of the prenyltransferase hexamer of PvCPS and a molecular model of the class II cyclase domain were used to interpret a low resolution molecular envelope of the full-length hexamer calculated from small-angle X-ray scattering (SAXS) data, revealing a fascinating (α∼βγ)_6_ “starburst”-like architecture.^30^ Interestingly, the cryo-EM structure of the related prenyltransferase hexamer of PfCPS reveals a partially-open hexamer conformation thought to represent a possible intermediate between hexameric and octameric prenyltransferase assemblies.^31^

We previously reported the X-ray crystal structure of the prenyltransferase domain of S723T-PvCPS, showing IPP bound to the prenyltransferase domain in a nonproductive location.^32^ We hypothesized that if the IPP binding site corresponded to a possible isoprenoid diphosphate binding site as GGPP transits from the prenyltransferase domain to the cyclase domain, then a productive prenyltransferase-cyclase complex might be stabilized for visualization in cryo-EM studies of full-length S723T-PvCPS. However, while we do not observe prenyltransferase-cyclase association in this system, we do observe well-defined density for the class II cyclase domain. Accordingly, we now present the 2.9 Å-resolution cryo-EM structure of the cyclase domain in full-length S723T-PvCPS, confirming general features of the model of this domain previously fit to the molecular envelope derived from SAXS data.^30^ Due to its relatively small size, the 63-kD βγ cyclase domain would ordinarily be considered a challenging target for cryo-EM structure determination. Even so, cryo-EM data were of sufficient quality to yield an atomic resolution reconstruction. Additionally, we report biochemical assays showing that GGPP channeling from the prenyltransferase to the cyclase is operative in the (α∼βγ)_6_ hexamer of wild-type PvCPS.

## Methods

### Reagents

Unless overwise noted, chemicals and reagents used for protein expression and purification were purchased from Fisher Scientific, Millipore Sigma, or GoldBio and were used without further purification.

### Protein Preparation and Purification

**Wild-type PvCPS and S723T-PvCPS.** Codon-optimized plasmids of wild-type PvCPS (Uniprot: A0A348FUE1) and S723T-PvCPS were purchased from Genscript. All PvCPS constructs utilize a pET28a(+) vector with an N-terminal His tag and a TEV cleavage site for purification. Detailed methods for the purification of PvCPS including wild-type and mutants have been previously reported and were followed without major modifications.^30^ Briefly, BL21(DE3) competent *E. coli* cells (New England Biolabs) were transformed with PvCPS plasmids and grown on agar plates. Six 2-L flasks containing 1 L of terrific broth (TB) supplemented with 50 μg/mL kanamycin were inoculated using 15-mL aliquots of overnight starter cultures. Expansion cultures were shaken at 37°C and 250 RPM until the OD reached approximately 0.6, at which point isopropyl β-D-thiogalactopyranoside was added to a final concentration of 500 μM. After shaking at 16°C for 18 h, cells were pelleted by centrifugation at 7808*g* for 20 min. Cell pellets were collected and stored at −80°C until further use.

Upon thawing, the cell pellet was resuspended in 100 mL Buffer A [25 mM 4-(2-hydroxyethyl)-1-piperazineethanesulfonic acid (HEPES) (pH 7.5), 250 mM NaCl, 30 mM imidazole, 1 mM tris(2-carboxyethyl)phosphine (TCEP), and 10% glycerol] also containing 100 mg lysozyme, 2 EDTA-free cOmplete Mini Protease Inhibitor tablets, 250 units Benzonase, and approximately 17 mg of phenylmethylsulfonyl fluoride (PMSF; dissolved in 100% ethanol prior to addition to the lysate mixture). The lysate was stirred at room temperature for 45 min and at 4°C for 30 min followed by sonication (30 Hz, cycling between 1 s on/2 s off for 10 min). The lysate mixture was clarified by centrifugation at 41,646*g* for 30 min at 4°C. The supernatant was loaded onto a 5-mL HisTrap HP prepacked column (Cytiva) pre-equilibrated with Buffer A. Protein was eluted with an initial 10% step to Buffer B [Buffer A plus 500 mM imidazole], followed by a 50 mL gradient to 100% Buffer B. Fractions containing the largest quantities of PvCPS were pooled, filtered with a 0.22 μM PVDF syringe filter (Millipore), and loaded onto a 26/60 Superdex 200 size-exclusion column (Cytiva) pre-equilibrated with sizing buffer [25 mM HEPES (pH 7.5), 150 mM NaCl, 10% glycerol, 1 mM TCEP]. Fractions were analyzed using SDS-PAGE and pure fractions were pooled and supplemented with 100% glycerol to bring the final glycerol concentration to 30%. The sample was then concentrated using a 50 kD Amicon centrifugal filter unit to 10 mg/mL and stored at −80°C.

**PvCPS_CY_ and PvCPS_PT_.** Purification of wild-type PvCPS was achieved as described above with the addition of a 1-h incubation with Endoproteinase Glu-C (Hampton Research) at a 1:1000 mg ratio at room temperature. The mixture was then reloaded onto a 26/60 Superdex 200 size-exclusion column (Cytiva) pre-equilibrated with sizing buffer [25 mM HEPES (pH 7.5), 150 mM NaCl, 10% glycerol, 1 mM TCEP]. Two major peaks eluted, the first containing an approximately 37-kD protein and the second containing an approximately 63-kD protein based on SDS-PAGE analysis. Fractions associated with the first and second peak were pooled as PvCPS_PT_ and PvCPS_CY_, respectively. PvCPS_PT_ was concentrated to 10 mg/mL, PvCPS_CY_ was concentrated to 20 mg/mL, and both were stored at −80°C.

**PvCPS_SL_.** A construct of PvCPS was designed in which the 125-residue linker segment L567–T692 was replaced by the 20-residue segment AISDHTSRAIDLCRNPPLWG corresponding to an extension of helix A in the prenyltransferase domain of PfCPS. The codon-optimized plasmid was purchased from Genscript. The expression and purification of this variant containing a shortened linker, designated “PvCPS_SL_”, were carried out in the same manner as described above for full-length PvCPS.

**CotB2.** A pET28a(+) expression plasmid containing the gene encoding cyclooctatenol synthase (CotB2) was purchased from Genscript. Previously published protocols^33,34^ were utilized to express and purify the single domain class I cyclase CotB2 as described without modification (Uniprot ID: C9K1X5).

**PaFS_CY_.** The cyclase domain of PaFS, designated “PaFS_CY_”, consists of residues 1–344.

The preparation of this construct was achieved using previously reported procedures without modification.^26^

### Mass Photometry

Samples of wild-type PvCPS and PvCPS_SL_ were initially diluted to approximately 50 nM in mass photometry (MP) buffer [25 mM HEPES (pH 7.5), 250 mM NaCl]. The mass photometer (Refeyn Ltd.) was calibrated using a standard curve based on measurements with β-amylase and thyroglobulin, which allowed for the conversion of ratiometric contrast to particle mass in kD. A typical measurement involved addition of 12 µL of mass photometry buffer to the glass slip receptacle which was first utilized to obtain optimal focus. Following this, 3 µL of the diluted protein sample was mixed with the drop to yield a final protein concentration of approximately 10 nM. Data were collected for 60 s (approximately 60,000 frames) and analyzed using RefeynMP software.

### Cryo-EM Data Collection

S723T-PvCPS was utilized for cryo-EM structure determination. An 80-μL sample (3.0 mg/mL) was dialyzed overnight in cryo-EM buffer (25 mM HEPES pH 7.5, 50 mM NaCl, 1 mM TCEP) and then filtered using a 0.22 μM PVDF centrifugal filter (Millipore) to yield a concentration of approximately 2.0 mg/mL. A 10X ligand and detergent stock was made containing MgCl_2_, IPP, and NP-40 detergent mixed in cryo-EM buffer and filtered with a 0.22 μM polyvinylidene fluoride (PVDF) centrifugal filter (Millipore). Protein was diluted to 1.0 mg/mL (approximately 10 μM) and the substrate mix was added to final concentrations of 5 mM MgCl_2_, 0.5 mM IPP, and 0.025% NP-40. Grid setup utilized R1.2/1.3 300-mesh copper grids (QUANTIFOIL™) which were glow discharged for 1 min (easiGlow, Pelco) prior to blotting. A Vitrobot Mark IV was utilized to blot 3 μL of protein sample with a blot force of 0 for 6 s prior to vitrification in ethane. Frozen grids were clipped and transferred to a Titan Krios G3i cryogenic transmission electron microscope (Thermo Fischer Scientific) operating at 300 keV at the Laboratory for Biomolecular Structure at Brookhaven National Laboratory (Upton, NY). Images were recorded with a K3 Summit detector at 81,000x magnification (0.53 Å/pixel) with a defocus of −0.8 to −2.5 µM. Data were collected utilizing EPU software with parameters of 40 frames taken at a dose of 15 e^−^/pixel/s (33 e^−^/Å^2^). A total of 3,310 movies were recorded.

### Cryo-EM Data Processing

Data processing was completed using cryoSPARC (version 4.6.2), summarized by the workflow shown in Figure S1. Movies were imported into cryoSPARC and aligned using patch motion correction with output Fourier cropping set to 1/2. Contrast transfer function (CTF) estimations were calculated with patch CTF estimation and resulting exposures were manually curated resulting in the rejection of 87 micrographs. An initial selection of particles was obtained by using the Blob Picker in cryoSPARC with a minimum and maximum particle diameter of 150 Å and 300 Å, respectively. The resulting particle picks were extracted and utilized to establish an initial set of two-dimensional (2D) classes. The best classes were selected and included in a template picker job in cryoSPARC set to identify particles with a diameter of 200 Å; following inspection, this procedure yielded 1,686,323 particles. Extraction of particles with a large box size (400 pixels), which would accommodate a hexameric particle, revealed low quality 2D classes of almost exclusively βγ domain monomers. To generate higher quality 2D classes of βγ domain monomers, particles were extracted with a smaller box size of 180 pixels Fourier-cropped to 90 pixels and utilized to establish 200 2D classes. Optimal particles were selected and utilized for *ab initio* three-dimensional (3D) reconstruction with two classes. The best volume was lowpass filtered to 20 Å and utilized for several rounds of heterogeneous refinement for both classes. The resulting highest resolution class was utilized for non-uniform refinement. Despite promising 2D classes of the class II cyclase, the final 3D reconstruction in cryoSPARC still suffered greatly from alignment issues, reflecting the significant challenge of working with a relatively small 63-kD particle. This particle size approached the lower limit of current cryo-EM capabilities (Figure S1).

A new cryo-EM tool in RELION, Blush regularization, has recently been shown to yield high-resolution 3D reconstructions of low molecular weight biomolecules using data with low signal-to-noise ratio; for example, application of Blush regularization yielded a 2.5 Å-resolution map of a 40-kD protein-RNA complex.^35^ To obtain a high-resolution 3D reconstruction of the βγ domains of S723T-PvCPS, the highest quality 2D classes were selected for a stack of approximately 500,000 particles. CTF-corrected micrographs with these particle picks were transferred from cryoSPARC to RELION, and particles were re-extracted with a box size of 180 pixels. 3D refinement was run with all 500,000 particles followed by 3D classification. Optimal 3D classes were used to select a subset of particles for a final stack of approximately 350,000 particles. This particle stack was subject to iterative rounds of 3D refinement, post-processing for B-factor sharpening, and CTF refinement to establish a set of polished particles. These particles yielded a final 3D reconstruction at 2.9 Å-resolution as indicated by the Gold Standard Fourier Shell Correlation plot (Figure S2).

A model of the S723T-PvCPS βγ cyclase domain was generated with AlphaFold3^36^ and fit into the 2.9 Å-resolution map using ChimeraX.^37^ The fit model and map were then transferred to PHENIX for real-space refinement using secondary structure restraints and simulated annealing to optimize the model fit to the experimental map.^38^ Iterative cycles of model adjustment with Coot^39^ and real-space refinement yielded the final model of the class II cyclase validated with the cryo-EM validation tool in PHENIX as well as with Molprobity.^40^ Data collection and refinement statistics are recorded in Table S1.

### Substrate Competition Experiments

Wild-type PvCPS and PvCPS_SL_ were utilized for substrate competition experiments designed to probe GGPP channeling between prenyltransferase and cyclase domains. Assays were performed on a 200-μL scale in 2-mL glass screw-cap vials. Reactions were run in sizing buffer also containing 1 mM MgCl_2_ in which the concentrations of PvCPS and diterpene cyclase competitor (CotB2 or PaFS_CY_) were 5 μM (MgCl_2_ concentrations higher than 1 mM were found to be significantly inhibitory for the generation of copalyl diphosphate/copalol). Experiments were also conducted with equimolar 5 μM mixtures of individual PvCPS prenyltransferase (PvCPS_PT_), PvCPS cyclase (PvCPS_CY_), and diterpene cyclase competitors mentioned above. A mixture of IPP and DMAPP was prepared by adding equal volumes of 10 mM DMAPP and 30 mM IPP. Reactions were initiated by the addition of 20 μL of either the DMAPP+IPP substrate mixture or a 5 mM solution of GGPP. All isoprenoids were purchased from isoprenoids.com and were resuspended in a 7:3 methanol:10 mM NH_4_HCO_3_ mixture at various concentrations.

Reactions were incubated at room temperature overnight. The following day, rCutSmart Buffer (New England Biolabs) was added to each reaction to a final concentration of 1X from a 10X stock, followed by the addition of 40 units of Quick CIP (New England Biolabs). Reaction mixtures were then incubated at 37°C overnight. The following day, reactions were quenched by addition of 200 μL ethyl acetate and then vortexed for 3 x 5-s intervals. The entire reaction volume was transferred to a 1.7-mL Eppendorf tube and spun at 2415*g* for 15 s. A portion of the organic phase was subsequently transferred to a screw cap vial with a 300-μL insert.

Gas chromatography-mass spectrometry (GC-MS) was employed for product identification and quantification using an Agilent 8890 GC / 5597C MSD system equipped with a J&W HP-5 ms Ultra Inert GC capillary column (30 m x 0.25 mm x 0.25 μm). While GC-MS can be utilized to detect and quantify cyclooctatenol, GC-MS cannot be utilized to detect and quantify copalyl diphosphate due to the presence of the diphosphate group. This necessitated a dephosphorylation step to generate copalol, which can then be extracted, detected, and quantified using GC-MS. Accordingly, dephosphorylation of copalyl diphosphate in product mixtures was achieved using calf intestinal phosphatase (CIP) as previously described by Ma and colleagues.^41^ To ensure complete dephosphorylation of copalyl diphosphate, CIP reactions were carried out for increasing time intervals until the copalol signal plateaued in control experiments. The optimal dephosphorylation time was determined to be 18 h.

Copalol-cyclooctatenol and copalol-fusicoccadiene product mixtures were analyzed using a GC program initiated with an oven temperature of 60°C held for 2 min followed by a ramp to 320°C at 10°C/min with a hold at 320°C for 2 min. Following a solvent delay of 3 min, EI+ mode was used to collect MS data. Diterpene products were identified and confirmed by comparison with spectra archived in the National Institute of Standards and Technology (NIST) database.

## Results

### Cryo-EM Structure

Cryo-EM grids of full-length S723T-PvCPS were prepared at a protein concentration of 1.0 mg/mL (approximately 10 μM), a concentration at which hexamers and monomers are observed by size exclusion chromatography-multi-angle light scattering (SEC-MALS).^30^ Mass photometry experiments with a 10^3^-fold lower protein concentration (10 nM PvCPS) indicate a mixture of oligomeric species, including monomer, dimer, trimer, tetramer, pentamer, and hexamer (Figure S3), consistent with the previously-reported concentration-dependence of PvCPS oligomerization.^30^ While oligomeric heterogeneity of PvCPS persists at the 10 μM concentration used to prepare cryo-EM grids, prenyltransferase hexamers were visible in 2D class averages (Figure 2A). Because a relatively small number of prenyltransferase hexamer particles were observed, and because we previously reported high-resolution X-ray crystal structures of wild-type and S723T-PvCPS prenyltransferase hexamers,^30,32^ we did not pursue a 3D reconstruction of the prenyltransferase hexamer. Finally, mass photometry experiments indicate that 10 nM PvCPS_SL_ is predominantly hexameric (Figure S3), but this variant did not yield suitable grids for cryo-EM analysis.

**Figure 2.**
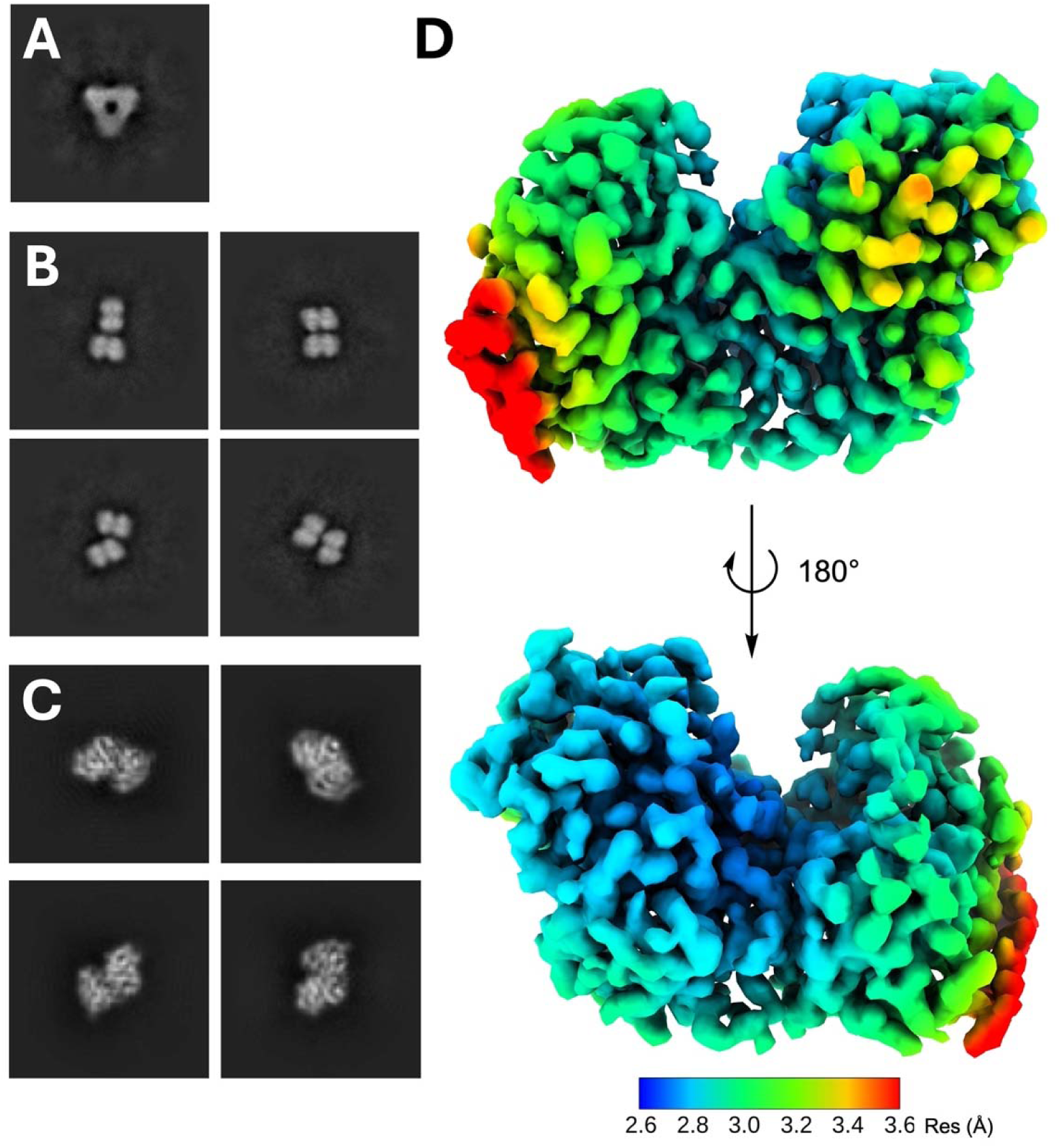
Representative 2D classes and final 3D reconstruction of the class II cyclase domain. (A) 2D class showing a hexameric assembly corresponding to the prenyltransferase domain. (B) 2D class averages showing βγ-βγ pairs. (C) High quality 2D classes of individual βγ domains. (D) Final 3D reconstruction of the class II cyclase with local resolution estimation as indicated.

Associated cyclase domains were not visible in 2D class averages of S723T-PvCPS prenyltransferase hexamers, consistent with a model in which cyclase domains are randomly splayed out from the prenyltransferase domains regardless of the oligomeric state of the protein. This is consistent with our previously reported cryo-EM studies of copalyl diphosphate synthase from *Penicillium fellutanum* (PfCPS).^31^ Despite the oligomeric heterogeneity of S723T-PvCPS in cryo-EM grids, reconstructions of the class II cyclase domain would be insensitive to oligomeric state. Since the S723T substitution occurs in the prenyltransferase domain, these cryo-EM data provided an ideal opportunity for closer study of the wild-type cyclase domain. Indeed, these cryo-EM data yielded 2D class averages of the 63-kD class II cyclase that clearly revealed secondary structure and characteristic βγ domain architecture. Interestingly, some βγ didomain particles were paired in close proximity with others in various orientations, suggesting the possibility of weak βγ-βγ interdomain interactions (Figure 2B).

Although 2D classes of the class II cyclase appeared to be promising (Figure 2C), achieving a high-resolution 3D reconstruction remained a significant challenge in view of the relatively small size of the particle, which at 63-kD approaches the lower limit of current cryo-EM capabilities. However, deployment of a newly developed small-particle processing pipeline in RELION, Blush regularization, enabled a 3D reconstruction at 2.9 Å-resolution. This reconstruction thus serves as a noteworthy example of cryo-EM structure determination of a relatively small protein. The reconstruction clearly reveals the expected didomain architecture of βγ domain assembly. The final map reveals all 12 helices of the β domain; however, two helices of the γ domain are characterized by ambiguous and weak density, so only 9 helices are included in the final model (Figure 2D).

The final cryo-EM reconstruction of the PvCPS class II cyclase domain allows for visualization of active site residues (Figure 3A, 3B) and can be compared with other class II cyclases that generate a variety of copalyl diphosphate stereoisomers or regioisomers (Figure 3C, 3D). To date, structures of the following copalyl diphosphate synthases are available in the Protein Data Bank (PDB): (+)-copalyl diphosphate synthase contained in the bifunctional enzyme abietadiene synthase from *Abies grandis*,^42^ (+)-copalyl diphosphate synthase contained in the bifunctional enzyme miltiradiene synthase from *Selaginella moellendorffii*,^43^ *ent*-copalyl diphosphate synthase from *Arabidopsis thaliana*,^20^ *ent*-copalyl diphosphate synthase from *Streptomyces platensis*,^44^ *syn*-copalyl diphosphate synthase from *Oryza sativa*,^41^ and 7,13-copalyl diphosphate synthase from *Grindelia robusta*.^45^ The structure of the PvCPS class II cyclase presented here is the first example of a fungal class II cyclase and the first example of a class II cyclase contained in a bifunctional prenyltransferase-cyclase system. PvCPS is also the third example of a class II cyclase that generates the copalyl diphosphate synthase stereoisomer shown in Figure 1, also known as (+)-or “normal” copalyl diphosphate.

**Figure 3.**
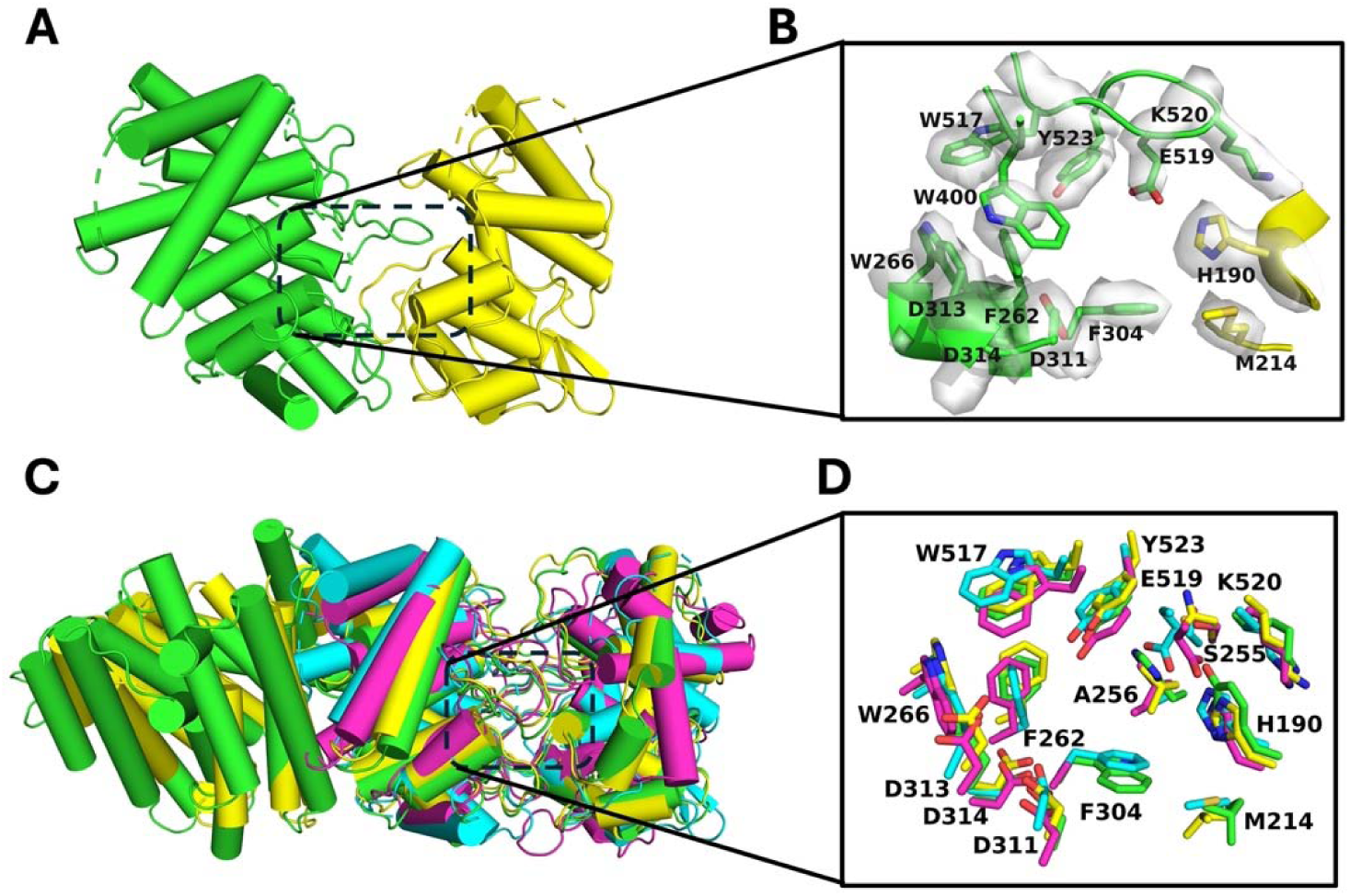
(A) Final model of the PvCPS class II cyclase (β domain = green, γ domain = yellow). (B) Active site of the class II cyclase domain showing the locations of key residues in the cryo-EM map, including general acid D313 in the DXDD motif along with several hydrophobic and aromatic residues. D313 is not characterized by well-defined density and side chain atoms are accordingly omitted from the final model shown. (C) Alignment of the βγ domains of PvCPS (cyan, PDB 9Q3I), a bacterial *ent*-copalyl diphosphate synthase (magenta, PDB 5BP8), copalyl diphosphate synthase from bifunctional abietadiene synthase (green, PDB 3S9V), and a plant *ent*-copalyl diphosphate synthase (yellow, PDB 4LIX) reveals generally similar overall protein folds. (D) Comparison of active site residues in aligned class II cyclases reveals generally similar active site structures, except for A256 and M214.

Comparisons of selected normal and *ent*-copalyl diphosphate synthases across different species reveal both conserved and subtly divergent structural features (Figure 3C, 3D). Alignment of the PvCPS cyclase domain with a bacterial *ent*-copalyl diphosphate synthase, a plant *ent*-copalyl diphosphate synthase, and the copalyl diphosphate synthase from bifunctional abietadiene synthase yields root-mean-square deviation (RMSD) values of 2.1, 3.8, and 3.4 Å, respectively, indicating generally similar overall structures. Several aromatic residues defining the active site contour, F262, W517, W266, and Y523, are conserved in these copalyl diphosphate synthases despite the fact that these enzymes generate two different product stereoisomers (Figure 3D). F262 and Y523 are substituted by Y317 and H501, respectively, in *syn*-CPS, where H501 serves as a catalytic base.^46^

Interestingly, certain active site residues differ slightly in their positioning or identity among the copalyl diphosphate synthases, including F304 and W400 in PvCPS (Figure 4). In PvCPS, W400 is positioned lower in the active site, potentially guiding the stereochemical outcome toward normal copalyl diphosphate. In *ent*- or *syn*-copalyl diphosphate synthases, the corresponding tryptophan is positioned higher in the active site (Figure 4), which may influence catalysis. Active site aromatic residues were recently subjected to mutagenesis in *ent*-CPS enzymes from *Erwinia tracheiphila* and *Arabidopsis thaliana*, revealing that the tryptophan equivalent to W400 in PvCPS is more important for catalytic efficiency rather than directing product outcome.^47^ The structures of CPS domains from abietadiene synthase from *Abies grandis* (AgAS), miltiradiene synthase from *Selaginella moellendorffii*, and CPS from *Grindelia robusta* are excluded from this comparison since the loop containing the corresponding tryptophan residue adopts an alternative conformation that does not substantially contribute to the active site contour (for example, PvCPS and AgAS are compared in Figure S4). Also of note, the diphosphate-binding region of copalyl diphosphate synthase active sites exhibits varying degrees of accessibility. In PvCPS and AtCPS this region is more constricted, whereas in *ent*-CPS and *syn*-CPS this region is much less constricted. The conserved glutamate in this region (E139 in PvCPS) coordinates to Mg^2+^ ions that are also coordinated by the substrate diphosphate group.^18,48^ Notably, the conserved glutamate in PvCPS is part of a “GFE” motif shared with other plant cyclases, and in bacterial cyclases it appears as G/AxE.^49^

**Figure 4.**
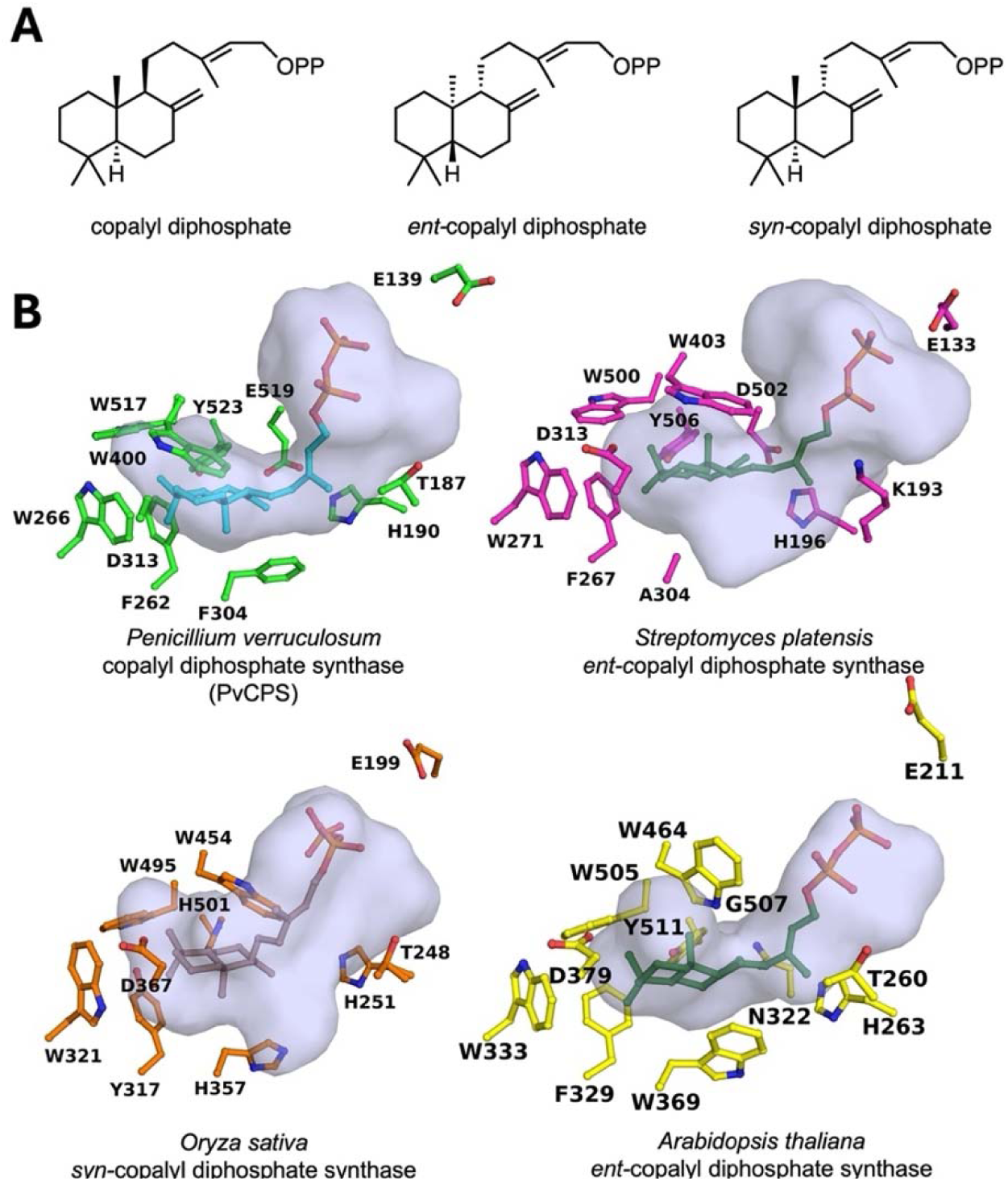
Product and active site structures of copalyl diphosphate synthases. (A) Molecular structures of copalyl diphosphate stereoisomers. (B) Active site contours of PvCPS, *ent*-CPS from *Streptomyces platensis*, *syn*-CPS from *Oryza sativa*, and *ent*-CPS from *Arabidopsis thaliana* reveal significant variations, some of which appear to be highly complementary to the particular product stereoisomer generated by the enzyme. The active site contour of PvCPS was generated from an AlphaFold3 model since some active site loops in the cryo-EM structure are disordered. However, residues shown for PvCPS derive from the cryo-EM structure.

A general base is required to quench the final carbocation intermediate. While this base is located in a conserved region of the active site, the identity of this general base varies depending on product stereochemistry and species of origin.^18,49–51^ Alternative general bases appear to guide the generation of alternative bicyclic diterpene skeletons. For example, tuberculosinyl diphosphate (TBP) synthase^52^ from *Mycobacterium tuberculosis* and terpentedienyl diphosphate (TPP) synthase^53^ from *Kitasatospora sp.* CB02891 generate halimane and clerodane diterpene skeletons, respectively, which differ from the labdane diterpene skeleton of copalyl diphosphate (Figure 5). The catalytic base in TBP synthase is a tyrosine positioned properly for a final deprotonation event accompanied by a series of 1,2-hydride and 1,2-methyl shifts to give the halimane diterpene product.^52^ In TPP synthase, the catalytic acid (D296) also serves as the catalytic base to yield the clerodane core.^53^

**Figure 5.**
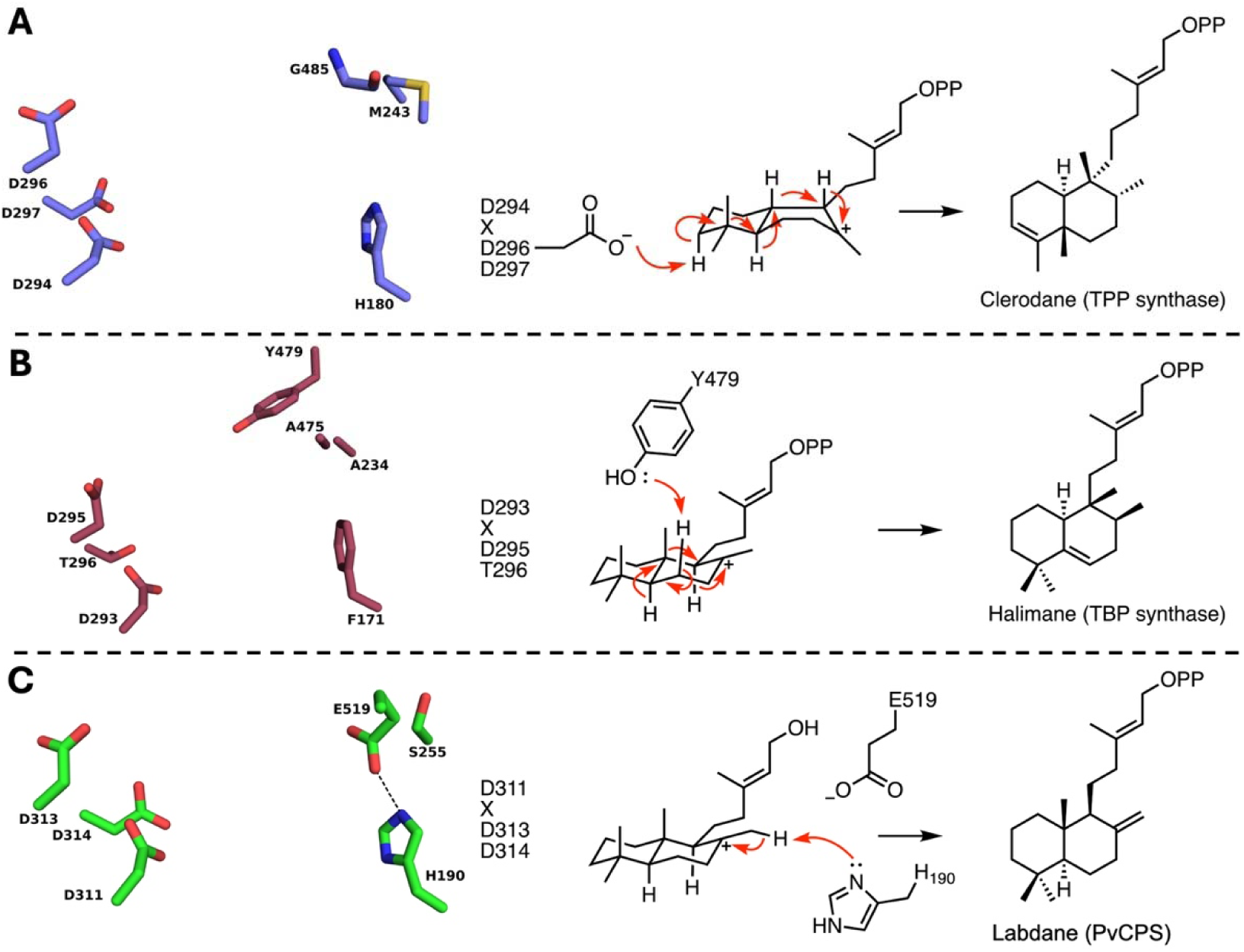
Molecular strategies for quenching the final carbocation intermediate in the generation of bicyclic diterpenes. (A) Terpentedienyl diphosphate (TPP) synthase is thought to utilize the conjugate base of general acid D296 to abstract an axial proton in the A ring, which is accompanied by four 1,2-methyl and -hydride migrations to yield the clerodane skeleton. While these migrations are illustrated as concerted for the sake of brevity, additional carbocation intermediates enroute to the clerodane product are possible. (B) Tuberculosinyl diphosphate (TBP) synthase likely utilizes the phenolic side chain of Y479 as a general base to abstract a proton in the B ring, which triggers 1,2-hydride/1,2-methyl/1,2-hydride migrations to yield the halimane skeleton. Although shown as a concerted reaction, additional carbocation intermediates are possible enroute to the halimane product. (C) PvCPS may utilize the imidazole group of H190 or the carboxylate group of E519 as a general base to abstract a proton from the exo methyl group of the B ring to generate the labdane skeleton.

Examples of class II diterpene cyclases with product outcomes that differ only by stereochemistry, but which utilize distinct catalytic bases, have also been identified (Figure 6A–D). For example, it has been proposed that plant *ent*-CPS in gibberellin phytohormone biosynthetic pathways utilize a histidine-asparagine catalytic dyad contained in L**H**S and P**N**V motifs that positions a hydrogen bonded water molecule to serve as the general base that quenches the final carbocation intermediate.^18,50,51,54,55^ However, these motifs are not otherwise conserved and also include histidine-tyrosine or histidine-threonine pairs (the latter involving just the first motif as L**HT**).^18, 56–58^ Interestingly, LHS and PNV motifs are conserved in CPS enzymes involved in gibberellin biosynthesis in bacteria, but not in fungi.^51^ Mutagenesis studies of *ent*-CPS from the fungus *Gibberella fujikuroi* indicate a conserved function for the histidine in the L**H**S motif as well as the threonine in an SG**T** motif that replaces the P**N**V motif in this enzyme.^51^ The histidine-threonine dyad is thought to mediate the final deprotonation step leading to *ent*-copalyl diphosphate formation. The crystal structure of *Streptomyces platensis ent*-CPS synthase reveals a unique aspartate-histidine dyad in which the aspartate is contained in a different loop compared to the asparagine in the P**N**V motif; even so, this dyad is proposed to activate a hydrogen bonded water molecule or utilize the aspartate-activated histidine to serve as a general base.^44,51^ Sequence alignment of various CPS enzymes illustrates the diversity of catalytic dyads that appear to serve general base functions (Figure 6E).

**Figure 6.**
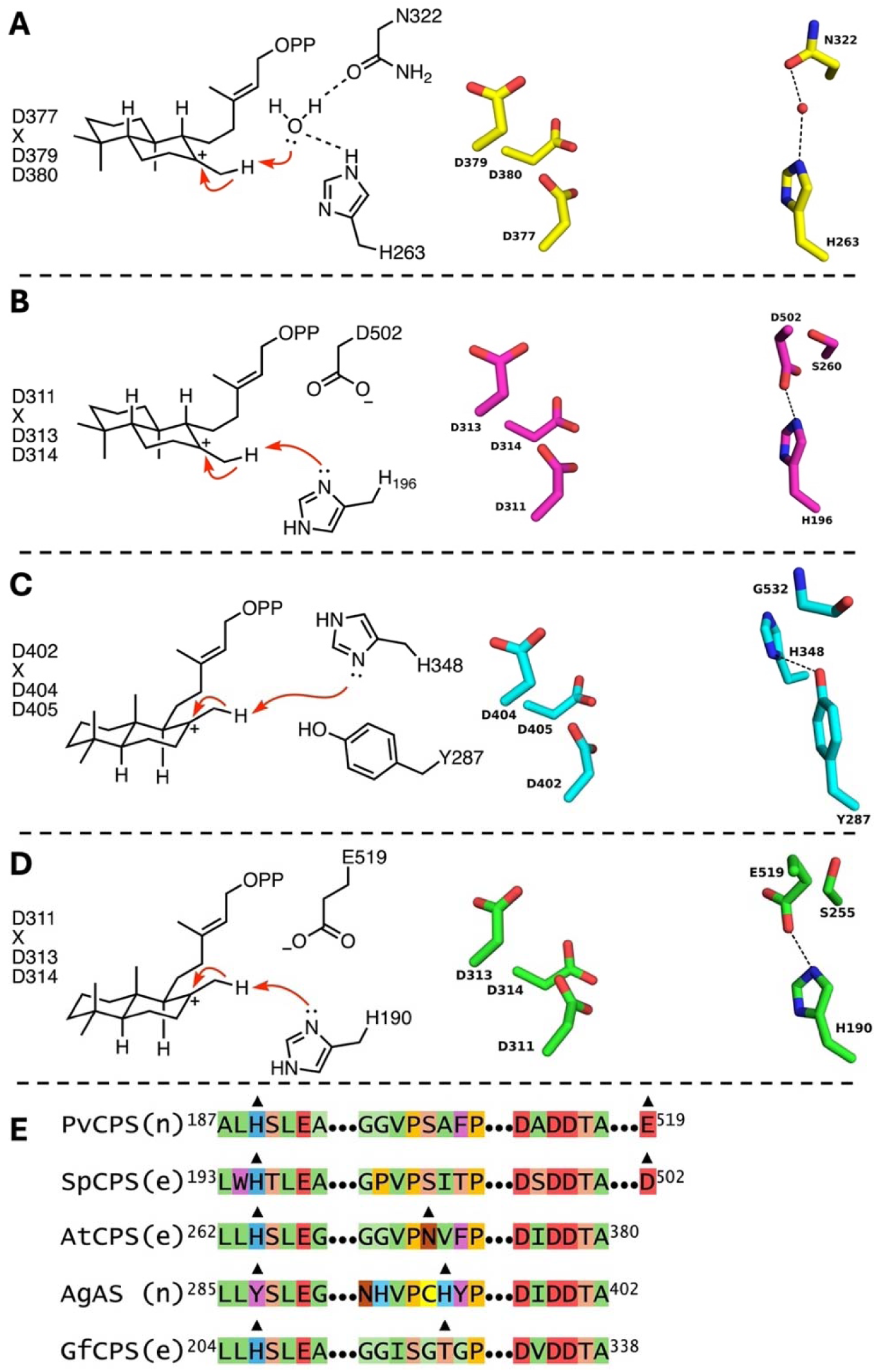
Catalytic dyads thought to serve general base functions in the generation of normal, *ent*-, and *syn*-copalyl diphosphate stereoisomers. (A) The catalytic dyad in *ent*-CPS from *Arabidopsis thaliana* consists of an asparagine and histidine that hydrogen bond to a water molecule thought to mediate the final deprotonation. (B) The catalytic dyad in *ent*-CPS from *Streptomyces platensis* consists of an aspartate and a histidine. (C) The catalytic dyad in *Abies grandis* abietadiene synthase (AgAS) consists of a histidine-tyrosine pair. (D) The catalytic dyad of PvCPS consists of a glutamate and histidine, the latter of which may serve to mediate the final deprotonation step. (E) Sequence alignment of various copalyl diphosphate synthases, indicating product stereochemistry as (n) for normal and (e) for *ent*. Triangles indicate residues comprising the catalytic dyad in each enzyme: PvCPS, CPS from *Penicillium verruculosum*; SpCPS, *ent*-CPS from *Streptomyces platensis*; AtCPS, *ent*-CPS from *Arabidopsis thaliana*; AgAS, CPS contained in abietadiene synthase from *Abies grandis*; GfCPS, *ent*-CPS from *Gibberella fujikuroi*.

### Substrate Channeling

Recent studies show that certain bifunctional class I terpene synthases are capable of channeling GGPP directly from the prenyltransferase to the cyclase.^25–27^ It has not yet been determined whether a bifunctional class II terpene synthase can similarly engage in substrate channeling. Cryo-EM studies of bifunctional class I terpene synthases suggest that weak, transient association of a cyclase domain with the side of a prenyltransferase oligomer facilitates substrate channeling.^26–29^ However, cryo-EM studies of PvCPS reported herein do not reveal any particles showing prenyltransferase hexamers with side-on binding of cyclase domains.

To ascertain the possibility of substrate channeling in PvCPS, we performed a competition assay in which an equimolar mixture of PvCPS and the diterpene cyclase cyclooctatenol synthase (CotB2) or fusicoccadiene synthase (PaFS) were incubated with GGPP. In a second experiment, the equimolar enzyme mixture was incubated with DMAPP and IPP such that the only source of GGPP in the reaction mixture was that generated by the PvCPS prenyltransferase. If substrate channeling were operative, the ratio of copalyl diphosphate-derived copalol/cyclooctatenol or copalol/fusicoccadiene would increase when the enzyme mixture is incubated with DMAPP and IPP, indicating that GGPP generated by the prenyltransferase remains on the enzyme for cyclization to form copalyl diphosphate.

Quantitation of copalyl diphosphate-derived copalol and cyclooctatenol using GC-MS reveals that the copalol/cyclooctatenol ratio increases, by more than five-fold, when the enzyme mixture is incubated with DMAPP and IPP compared to GGPP (Figure 7, Table 1). If the PvCPS cyclase and prenyltransferase are included as separate constructs along with the competitor cyclase, the ratio with DMAPP and IPP still increases, but to slightly less than five-fold. When an equimolar mixture of PvCPS_SL_ and CotB2 is studied using the same competition assay, the copalol/cyclooctatenol ratio increases by more than three-fold when the enzyme mixture is incubated with DMAPP and IPP compared to GGPP (Figure 7, Table 1). These observations suggest that much but not all of the GGPP generated by the PvCPS prenyltransferase remains on or close to the enzyme for cyclization to form copalyl diphosphate, i.e., there is some degree of substrate channeling from the prenyltransferase to the cyclase regardless of whether they are covalently linked or not. Channeling is not completely efficient, however, since GGPP is also released and accessible to cyclase competitors. These results are consistent with a model in which transient prenyltransferase-cyclase association facilitates GGPP transfer due to proximity of active sites, but the optimal positioning of prenyltransferase and cyclase domains is unknown. Channeling appears to be slightly less effective in PvCPS_SL_, perhaps because cyclase domains are held in a less favorable position relative to the prenyltransferase due to the shorter linker.

**Figure 7.**
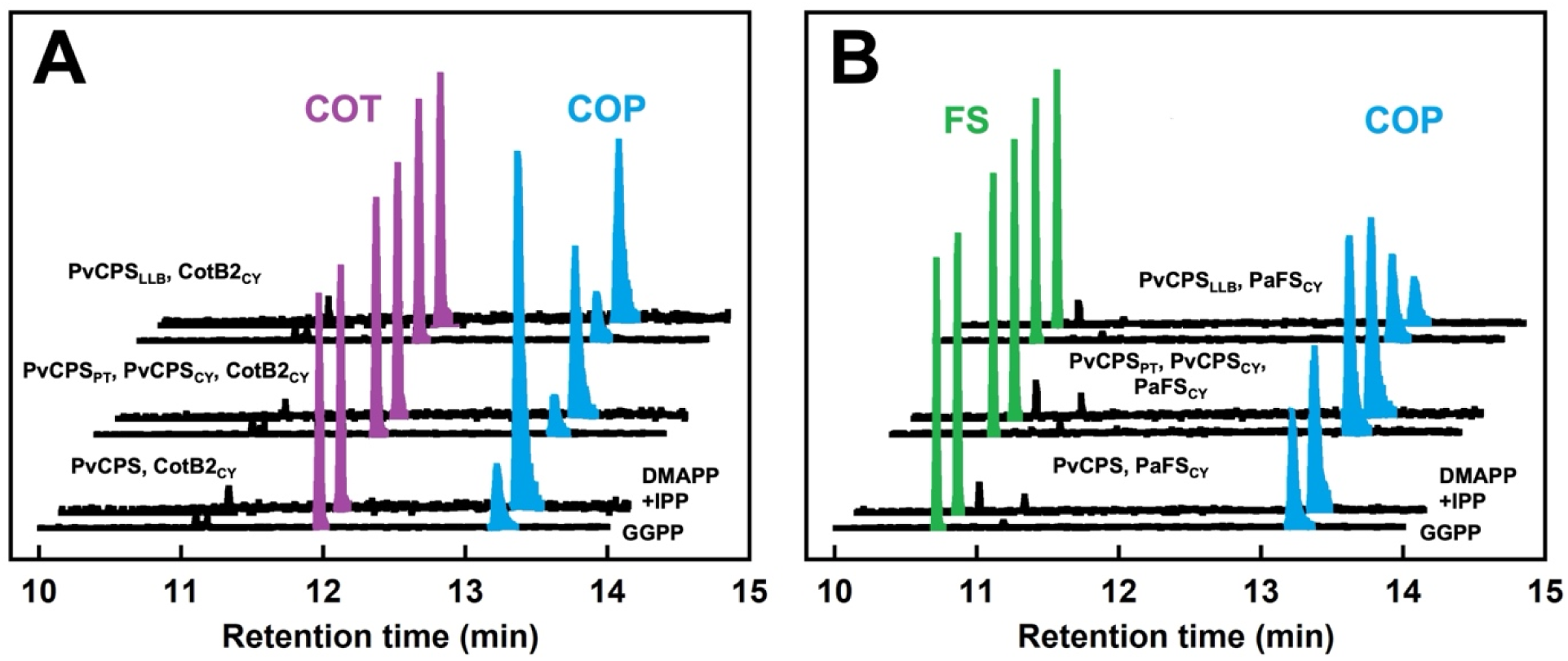
Products resulting from substrate competition experiments with PvCPS or CotB2 (A) and PvCPS and PaFS_CY_ (B). GC traces reveal products resulting from incubation of the equimolar enzyme mixture with GGPP (front trace) or DMAPP and IPP (back trace). COT, cyclooctatenol; COP, copalol; FS, fusicoccadiene. Traces are normalized such that the largest peak is equivalent in size.

**Table 1.** Product ratios resulting from substrate competition experiments.

| Equimolar Enzyme Mixture | Products Formed<sup>a</sup> | Product Ratio from GGPP | Product Ratio from DMAPP and IPP |
| --- | --- | --- | --- |
| PvCPS, CotB2 | COP:COT | 0.64 ± 0.08 : 1 | 3.33 ± 0.18 : 1 |
| PvCPS, PaFS<sub>CY</sub> | COP:FS | 1.06 ± 0.07 : 1 | 1.55 ± 0.13 : 1 |
| PvCPS<sub>PT</sub>, PvCPS<sub>CY</sub>, CotB2 | COP:COT | 0.29 ± 0.03 : 1 | 1.35 ± 0.03 : 1 |
| PvCPS<sub>PT</sub>, PvCPS<sub>CY</sub>, PaFS<sub>CY</sub> | COP:FS | 1.98 ± 0.16 : 1 | 2.26 ± 0.23 : 1 |
| PvCPS<sub>SL</sub>, CotB2 | COP:COT | 0.43 ± 0.05 : 1 | 1.41 ± 0.10 : 1 |
| PvCPS<sub>SL</sub>, PaFS<sub>CY</sub> | COP:FS | 1.05 ± 0.10 : 1 | 0.52 ± 0.01 : 1 |

<sup>a</sup>COP, copalol; COT, cyclooctatenol; FS, fusicoccadiene

When the substrate competition experiment is performed with an equimolar mixture of PvCPS and PaFS_CY_, or an equimolar mixture of PvCPS_PT_, PvCPS_CY_, and PaFS_CY_, the copalol/fusicoccadiene ratio increases only slightly, by 1.46-fold, and 1.14-fold, respectively, when the enzyme mixture is incubated with DMAPP and IPP compared to GGPP (Figure 7, Table 1). Channeling in PvCPS appears to be marginal when the PvCPS cyclase competes with PaFS_CY_ for GGPP generated by the PvCPS prenyltransferase. Curiously, when the cyclase domain of PvCPS_SL_ competes with PaFS_CY_ for GGPP generated by the PvCPS_SL_ prenyltransferase domain, there is increased production of fusicoccadiene. This result suggests that GGPP preferentially transits to PaFS_CY_ rather than the cyclase domain of PvCPS_SL_. In other words, intermolecular channeling of GGPP to PaFS_CY_ appears to be preferred over intramolecular channeling of GGPP to the cyclase domain of PvCPS_SL_. Intermolecular GGPP channeling to PaFS_CY_ was similarly observed with variediene synthase, where a transient side-on binding mode of PaFS_CY_ to the prenyltransferase hexamer was proposed to account for intermolecular channeling.^27^ It is possible that a similar PaFS_CY_ binding mode with PvCPS_SL_ facilitates intermolecular GGPP channeling.

## Discussion

In the past two decades, cryo-EM has come to rival X-ray crystallography for protein structure determination at high resolution; indeed, proteins that are refractory to crystallization are often amenable to structure determination by cryo-EM.^59^ However, cryo-EM has its limitations as well, in that smaller proteins <100 kD often fail to yield satisfactory reconstructions due to alignment complications associated with a lower signal-to-noise ratio.^35^ To facilitate the study of smaller proteins using cryo-EM, enabling tools such as nanobodies have been developed to bind to a target particle so as to increase the particle size.^60^ Absent such approaches, high-resolution reconstructions of particles less than ∼100 kD are relatively rare, especially in cases where molecular symmetry cannot be exploited to enhance the signal-to-noise ratio. A hexamer of full-length PvCPS is approximately 660 kD and therefore should be a reasonable target for cryo-EM analysis. However, due to significant flexibility of the 125-residue linker connecting the α and βγ domains in each α∼βγ protomer, such that the six βγ domains of the class II cyclase are randomly splayed-out around a central prenyltransferase α_6_hexamer, high-resolution structure determination of the full-length protein using either cryo-EM or X-ray crystallography is impossible.

Even so, structure determinations of the individual catalytic domains of PvCPS are feasible. For example, crystallization of the prenyltransferase α_6_hexamer yielded a 2.41 Å-resolution X-ray crystal structure,^30^ crystal structures of site-specific prenyltransferase variants have been determined at resolutions ranging 2.00–3.15 Å,^32^ and the cryo-EM structure of the related prenyltransferase domain of PfCPS has been determined at 2.81 Å resolution.^31^ Although the βγ domain assembly of the PvCPS cyclase was not amenable to crystallization, we show here that it is amenable to atomic resolution structure determination by cryo-EM. Moreover, a separate construct was not required for structure determination, since cryo-EM grids prepared with full-length PvCPS yielded separate 2D reconstructions of the prenyltransferase hexamer and the class II cyclase domains. Since the structure of the prenyltransferase hexamer had already been reported, we focused our attention on the 63-kD class II cyclase domain. The use of the new Blush regularization tool in RELION^35^ enabled reconstruction of this relatively small particle at 2.9 Å-resolution.

The structure of the PvCPS cyclase domain is useful for advancing our understanding of structure-function relationships in the greater family of class II terpene cyclases. There are no structures reported to date for a class II cyclase from a fungal species, so it is instructive to compare the PvCPS cyclase with plant and bacterial class II cyclases. In particular, both bacterial and plant copalyl diphosphate synthases share a number of conserved residues lining the active site pocket, although key differences appear to direct alternative mechanistic pathways for the generation of alternative stereoisomers of the labdane skeleton. For example, plant enzymes involved in gibberellin biosynthesis have a histidine-asparagine catalytic dyad which interacts with a water molecule serving to facilitate the final deprotonation step leading to *ent*-CPS products.^18,50,51^ The identify of the catalytic base in class II diterpene cyclases has been found to be highly variable, though, with examples that include tyrosine-histidine or histidine-threonine dyads.^18, 56–58^ Notably, in a bacterial *ent*-CPS the catalytic dyad consists of an aspartate-histidine pair hydrogen bonded to a water molecule that may be responsible for the final proton abstraction.^44^ Interestingly, if G487 of a bacterial TPP synthase is substituted by aspartate, the product outcome is altered so as to generate a labdane skeleton instead of the native clerodane skeleton.^53^ The cryo-EM structure of the PvCPS cyclase reveals that E519 corresponds to the glutamate or aspartate observed in bacterial class II cyclases that generate the labdane core, suggesting that this residue similarly serves as a general base in catalysis. Since fungal species are more closely related to plants, it is interesting that PvCPS seems to have adopted the mechanistic strategy of a bacterial cyclase.

The current study is the first to reveal GGPP channeling from the prenyltransferase to the cyclase in a class II bifunctional terpene synthase. Transient prenyltransferase-cyclase association is proposed to account for intramolecular as well as intermolecular channeling, as previously proposed for certain class I bifunctional terpene synthases,^26–29^ but the mode of prenyltransferase-cyclase association in a class II system remains unknown. Of note, PvCPS oligomerization is sensitive to concentration, and oligomerization could conceivably influence GGPP channeling due to the increased number of enzyme active sites held in close proximity to each other. Previous biophysical studies of PvCPS suggest that this system does not form hexamers exclusively at 5 μM enzyme concentrations (the concentration utilized for channeling experiments), but instead is a mixture of hexamer and lower order oligomers.^30^ This is consistent with the relatively low occurrence of prenyltransferase hexamers found in 2D class averages of cryo-EM samples prepared with 10 μM enzyme concentrations (Figure 2A).

## Concluding Remarks

The 2.9 Å-resolution cryo-EM structure of the 63-kD class II cyclase domain of PvCPS is notable in that this relatively small protein yielded a reconstruction at atomic resolution. The structure of the PvCPS cyclase enables comparisons with other class II cyclases, including enzymes that generate alternative regioisomers or stereoisomers of copalyl diphosphate and related bicyclic diterpenes. Natural products derived from copalyl diphosphate, such as abietane and pimarane-type diterpenes, are found in the plant oleoresin secreted in response to insect attack or physical injuries.^61^ Copalyl diphosphate is also associated with molecules which have a broad range of medicinal applications, including modified grindelic acids and native sclareols that exhibit anti-tumor, anti-inflammatory, and anti-pathogenic activities.^62–64^ A greater understanding of the prenyltransferase-cyclase assembly line will advance synthetic biology approaches for the generation of valuable natural products. Future studies will probe the nature of transient prenyltransferase-cyclase association and the role of preferential GGPP transit in these systems.

## ASSOCIATED CONTENT

### Supporting Information

Supporting Information is available free of charge at:

Table S1, cryo-EM structure determination statistics; Figure S1, cryo-EM workflow; Figure S2, particle orientation and Gold-Standard Fourier Shell Correlation plot; Figure S3, mass photometry data; Figure S4, structural comparison of the W400 active site loop.

### Accession Codes

The cryo-EM map of the PvCPS cyclase has been deposited in the Electron Microscopy Data Bank (EMDB, www.ebi.ac.uk/pdbe/emdb) with accession codes EMD-72194. Atomic coordinates of the PfCPS prenyltransferase hexamer have been deposited in the Protein Data Bank (www.rcsb.org) with accession code 9Q3I.

## Funding

This research was supported by NIH grants R01 GM56838 to D.W.C. and R35 GM118090 to R.M. M.N.G. and K.S. were supported by Chemistry-Biology Interface NIH Training Grant T32 GM133398.

## Conflict of Interest Statement

The authors declare no competing interests.

## ACKNOWLEDGMENTS

This research utilized cryo-EM facilities at the Laboratory for BioMolecular Structure (LBMS), Brookhaven National Laboratory, which is supported by the DOE Office of Biological and Environmental Research (KP1607011).

## TOC Graphic

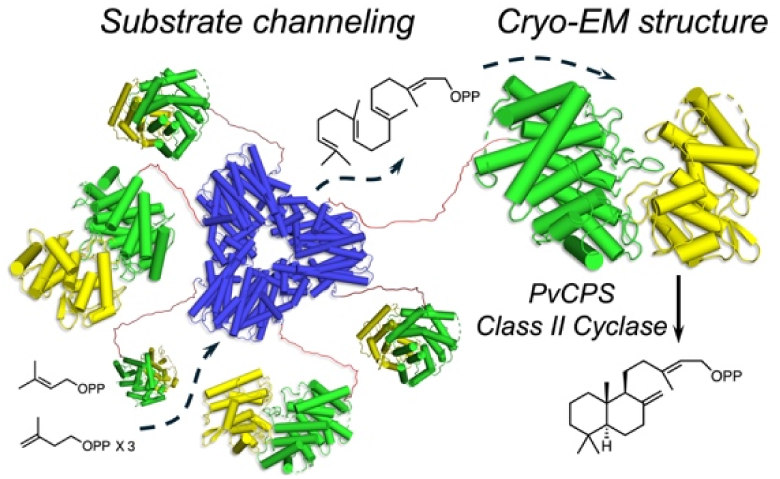

## Notes

### Competing Interest Statement

The authors have declared no competing interest.

### Summary of Updates

New coauthor added, minor text clariications, new citations added.

