## Supplementary Information for "Cryo-EM Structure of the Cyclase Domain and Evaluation of Substrate Channeling in a Bifunctional Class II Terpene Synthase"

**Table S1. Cryo-EM data collection parameters and refinement statistics**

| | PvCPS Cyclase (*C*1) |
|---|---|
| **Magnification** | 81000 |
| **Voltage (kV)** | 300 |
| **Exposure ($e^-$/Å$^2$)** | 33 |
| **Defocus range (μm)** | -0.8 to -2.5 |
| **Pixel size (Å/pix)** | 0.53 |
| **Symmetry imposed** | *C*1 |
| **Initial particles (no.)** | 1,686,323 |
| **Final particles (no.)** | 381,049 |
| **Map resolution (FSC = 0.143) (Å)** | 2.9 |
| **Model** | |
| ***Initial model used*** | AlphaFold3 |
| **Model composition (#)** | |
| **Chains** | 1 |
| **Atoms** | 3636 |
| **Residues** | 494 |
| **Water** | 0 |
| **Ligands** | 0 |
| **Root-mean-squared deviations** | |
| **Bond lengths (Å)**<br>**Bond angles (˚)** | 0.003<br>0.5 |
| **Validation** | |
| **MolProbity score** | 2.48 |
| **Clash score** | 13 |
| **Poor rotamers (%)** | 4.9 |
| **Ramachandran plot (%, MolProbity)** | |
| **Favored**<br>**Allowed**<br>**Outliers** | 95.36<br>4.01<br>0.63 |
| **Peptide plane (%)** | |
| **Cis proline/general**<br>**Twisted proline/general** | 7.7/0.0<br>0.0/0.0 |
| **B-factors (Å$^2$)** | |
| **(min/max/mean)** | 6/58/31 |
| **Model vs. Data** | |
| **CC (mask)**<br>**CC (box)**<br>**CC (peaks)**<br>**CC (volume)** | 0.62<br>0.67<br>0.64<br>0.64 |
| **PDB accession code** | **9Q3I** |
| **EMDB accession code** | **EMD-72194** |

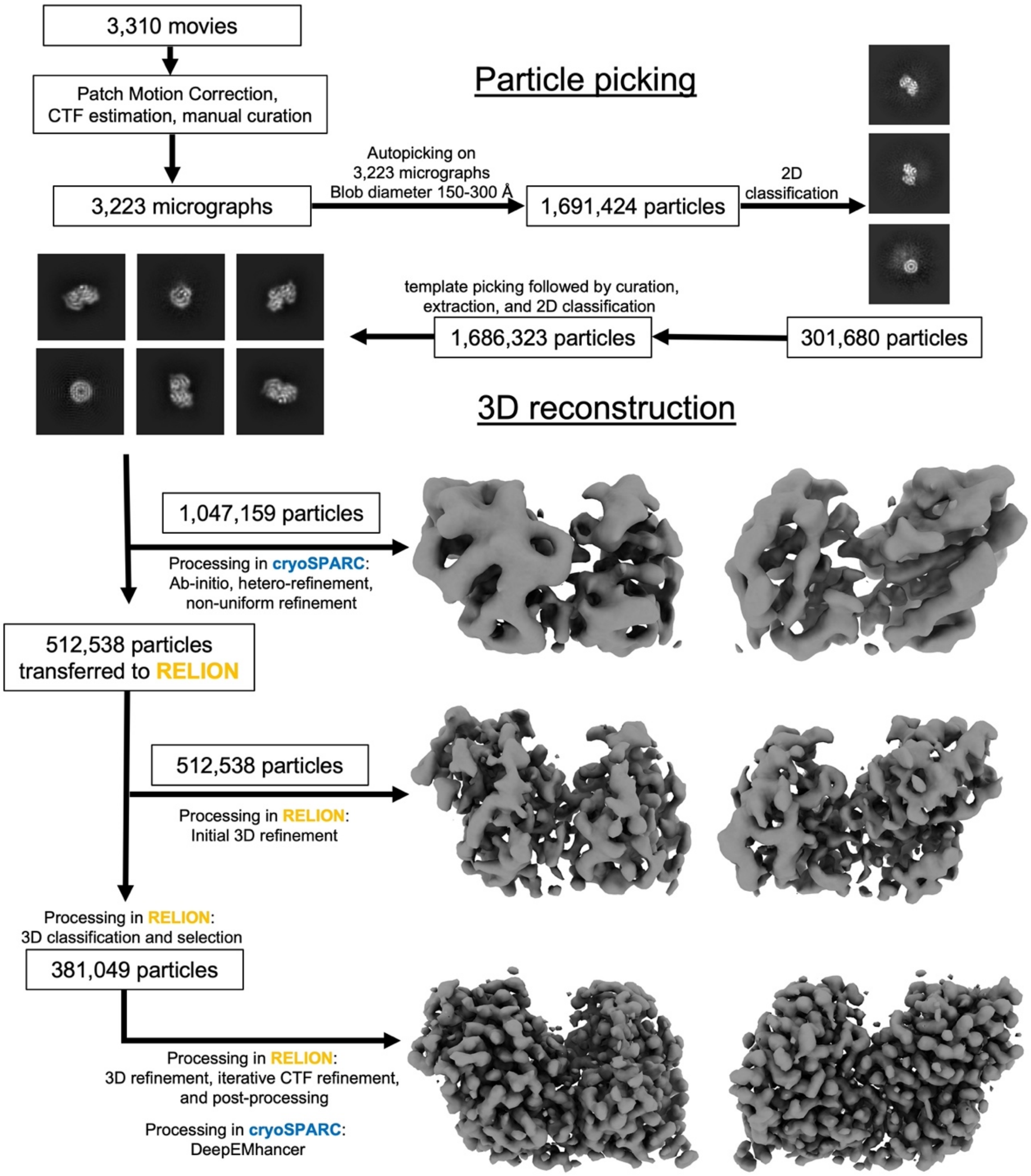


**Figure S1. Cryo-EM workflow for structure determination of the PvCPS class II cyclase.**

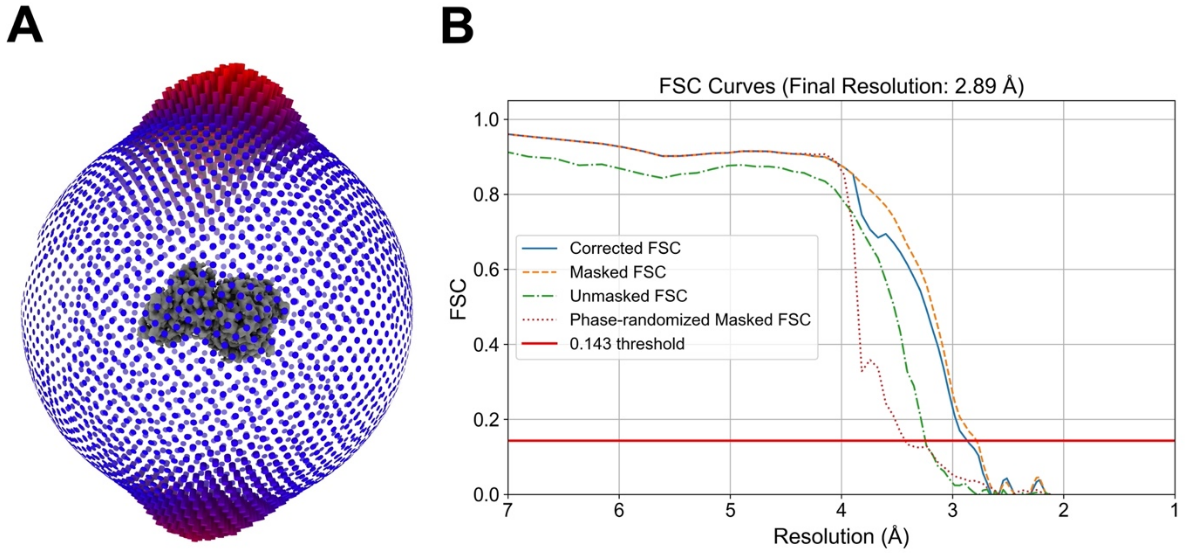


**Figure S2. Particle orientation distribution and GSFSC plot.** (A) Orientation distribution of particles utilized in the final reconstruction. (B) Gold-Standard Fourier Shell Correlation (GSFSC) curve following post-processing in RELION. The estimated resolution is 2.9 Å at GSFSC = 0.143.

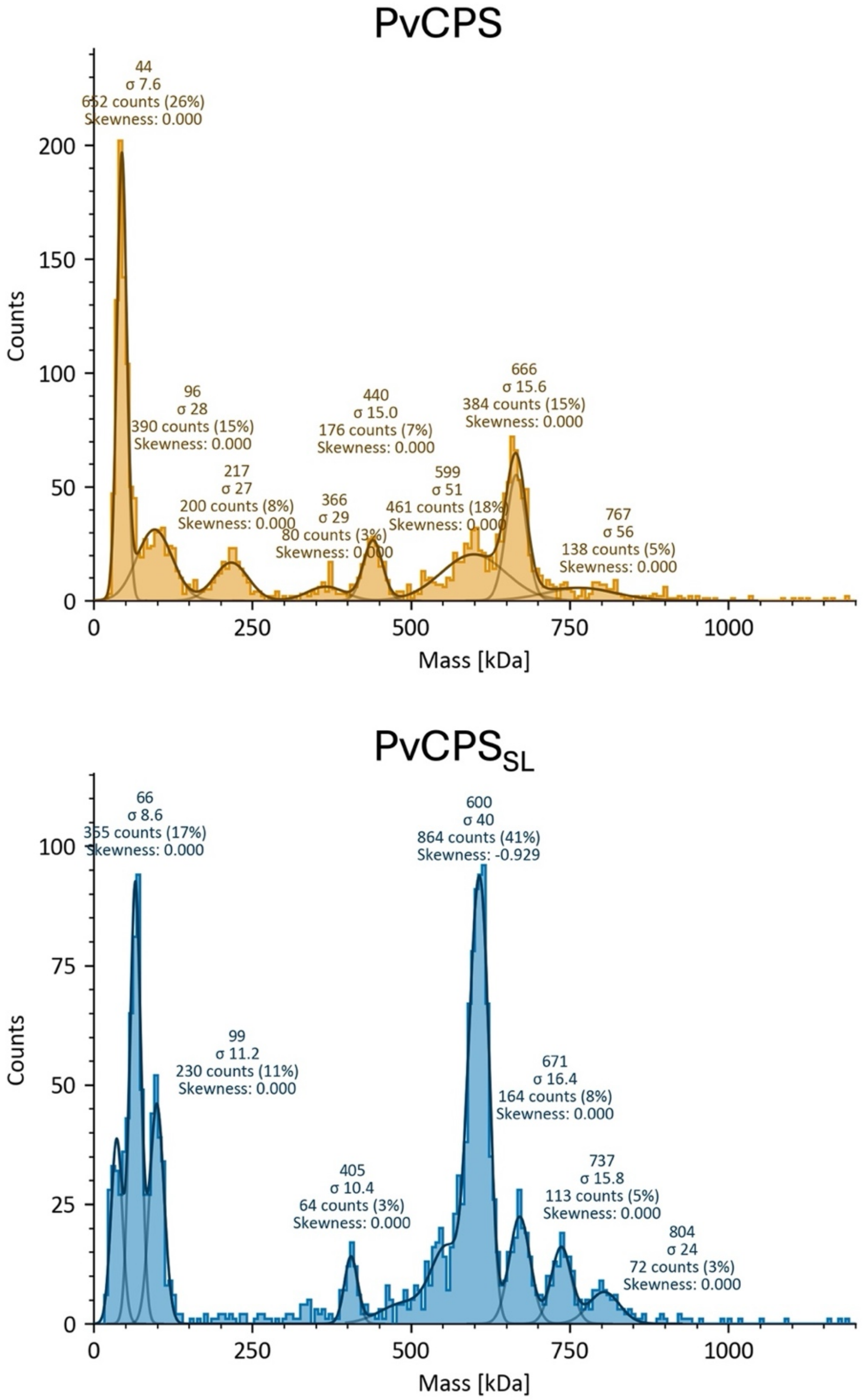


**Figure S3. PvCPS and PvCPS$_{SL}$ oligomers detected by mass photometry.** 10 nM PvCPS is a mixture of monomers, dimers, trimers, tetramers, pentamers, and hexamers, with hexamers being most prevalent. 10 nM PvCPS$_{SL}$ is predominantly hexameric, but monomers, tetramers, and octamers are also detected.

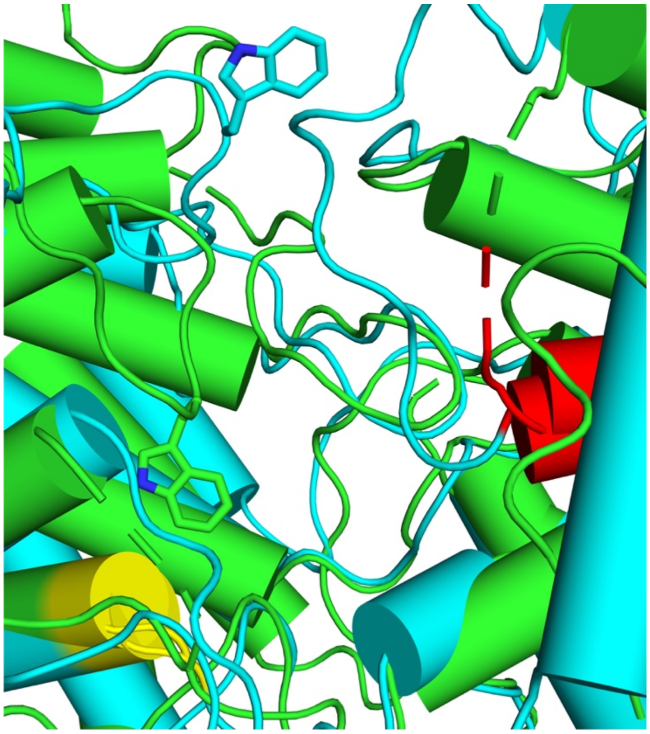

**Figure S4. Comparison of the W400 loop of PvCPS with the corresponding loop in the copalyl diphosphate synthase domain of abietadiene synthase from *Abies grandis* (AgAS).** Overlay of PvCPS (green) and AgAS (cyan). In PvCPS, W400 contributes substantially to the active site contour, but in AgAS the corresponding tryptophan is positioned such that it does not substantially contribute to the active site contour. For reference, the location of the aspartic acid that serves as a catalytic general acid is shown in yellow, and the location of the glutamate implicated in $Mg^{2+}$ complexation and hence substrate binding is shown in red.
